# Gαi2 coordinates neuronal microtubule and neurofilament networks to regulate axon initiation *in vivo*

**DOI:** 10.64898/2026.03.20.713162

**Authors:** Victoria E. Higgs, Danny Anwas, Artur Astapenka, Gabriela Toro-Tapia, Raman M. Das

## Abstract

Neurons possess a highly polarised morphology, established through axon formation. However, the mechanisms regulating stable axon initiation during embryonic development remain poorly understood. Here, using fixed-tissue super-resolution and high-resolution live-tissue imaging in developing chick spinal cord, we demonstrate that the multifunctional G-protein Gαi2 coordinates the interconnected neurofilament and microtubule networks to achieve stable axon outgrowth. Before axon initiation, microtubule network orientation shifts towards the site of the future axon where coiled Gαi2-associated neurofilaments accumulate asymmetrically before unfurling into the initiating axon behind microtubules. Crucially, Gαi2 is associated with neurofilaments and microtubules at points of contact. Gαi2 depletion reduces engagement between these cytoskeletal networks, and leads to impaired passage of neurofilaments into initiating axons and disrupted axon outgrowth, supporting a role for Gαi2 in regulating microtubule-driven incorporation of neurofilaments in nascent axons. These findings advance mechanistic understanding of polarity establishment and offer an enriched view of cytoskeletal regulation during axon formation.

## Introduction

Neuronal function relies on a highly polarised cellular morphology, which is established by the formation of a nascent axon during embryonic development (*1–3*). However, the mechanistic basis through which this process is regulated *in vivo* remains poorly understood. Embryonic central nervous system neurons differentiate from neuroepithelial progenitor cells, which possess apico-basal polarity and form the embryonic neuroepithelium (*4*). In the developing spinal cord, differentiating neurons undergo acute loss of polarity through abscission of the apical membrane (*5*). These newborn neurons then delaminate from the neuroepithelium and undergo repolarisation along an adjusted axis by *de-novo* protrusion of a nascent axon from the cell body (*C*) in a front–rear-type polarisation event. However, the cellular mechanisms facilitating this precise repolarisation are not well understood.

Cellular polarisation and axon outgrowth are dependent on coordinated asymmetric cytoskeletal remodelling (*1–3*). The early neuronal cytoskeleton consists of three major networks (*2*): actin, which forms dynamic filaments, generally at the cell periphery (*7*); microtubules, which are dynamic, strong polar filaments that provide directionality to polarising cells (*8*); and neurofilaments, which are long-lived, flexible intermediate filaments that provide structural support to the axon (*9*). Studies on the outgrowth of neurites (early axons or dendrites) once they have been initiated demonstrate that these nascent projections become engorged with microtubules and neurofilaments (*2, 10, 11*), which consolidate into longitudinal bundles to form the axon shaft (*3, 12*). At the distal neurite tip, a growth cone is established, with a microtubule-rich central region and a dynamic actin-rich periphery (*3*). However, little is known about the cytoskeletal dynamics that direct the initiation of axon outgrowth from a discrete region of the cell body. Axon outgrowth is preceded by accumulation of filamentous actin at the site of initiation (*2, 10, 13, 14*). Although it is known that this redistribution of actin is necessary for initiation of the protrusion (*10, 14*), the dynamics of microtubules and neurofilaments at this early stage, and the roles played by these networks in *de-novo* axon initiation are poorly understood. In dissociated hippocampal neurons cultured *in vitro*, microtubules have been observed to extend towards one side of the morphologically unpolarised cell (*15*) and preferentially grow into areas of the periphery that will form neurites (*16*). However, neurons cultured *in vitro* lack the tissue environment that directs axon formation (*17*). In addition, many of these cells are polarised postmitotic neurons at the point of plating, and existing protrusions are damaged and lost in the dissociation process, increasing the possibility that this model recapitulates repolarisation of cells that were already polarised *in vivo* and may therefore retain aspects of polarity (*18*). Consequently, the cytoskeletal dynamics of *de-novo* axon formation within the embryo are poorly understood.

The G protein Gαi2 is highly expressed in the developing chick spinal cord (*19*); however, its function in this context is unclear. Gαi proteins have diverse functions that depend on their subcellular localisation and interactions (*20*). In their canonical roles, Gαi proteins localise to cell membranes to transduce G-protein coupled receptor (GPCR) signalling via cAMP (cyclic AMP) as part of heterotrimeric G-protein complexes. Gαi proteins can also function independently of GPCRs and cAMP regulation in so-called non-canonical roles. Among these roles, which also include maintenance of Golgi structure , and steroid and tyrosine kinase receptor signalling (*20*), Gαi proteins have been shown to interact with and regulate the microtubule network. For example, Gαi facilitates positioning of the mitotic spindle through indirect interaction with astral microtubules via the LGN complex of proteins (*21*). In a recent study, Gαi2 was reported to regulate microtubule dynamics and cell polarity in migrating neural crest cells (*22*). In addition, related proteins Gαi1 and Gαs bind with tubulin *in vitro* (*23–25*) and are proposed to regulate cell shape by directly modulating microtubule dynamics (*26*). These studies suggest that Gαi proteins could have roles in regulating the cytoskeleton and cell polarity. Depletion of Gαi2 in the chick spinal cord results in a decrease in postmitotic neurons and an increase in early differentiating neurons (*1S*), suggesting disrupted neuronal differentiation. Additionally, Gαi2 depletion in mice leads to disrupted transition of neurons from a bipolar to a multipolar morphology in the cerebral cortex (*27*), suggesting that Gαi2 might regulate neuronal process outgrowth. Together, these studies suggest that Gαi2 has a role in regulating establishment of neuronal morphology during differentiation; however, the precise mechanistic basis of its function in this context is unclear.

Here, using a combination of fixed and live-tissue imaging of the developing chick spinal cord, we identify Gαi2 as a regulator of asymmetric cytoskeletal remodelling in *de-novo* axon formation. We report that the microtubule and neurofilament networks are polarised prior to axon initiation, with neurofilaments accumulated towards the site of axon initiation and centrosome-derived microtubules directed towards the accumulated neurofilaments. Gαi2 accumulates on the axon-directed neurofilament network, including at points of microtubule contact. Gαi2 knockdown reduces interaction between neurofilaments and microtubules, reduces the incorporation of neurofilaments into the initiating axon, and disrupts the precision and stability of axon formation. We propose that Gαi2 regulates the establishment and maintenance of neuronal polarity by mediating the microtubule-driven incorporation of stabilising neurofilaments into the nascent axon.

## Results

### Gαi2 accumulates towards the fronts of differentiating neurons before and during axon formation

To characterise the distribution of Gαi2 in the developing spinal cord, we fixed and immunolabelled E3 chick embryos and examined cross sections of spinal cord neural tube (Fig. 1 A, B). This revealed that Gαi2 protein was strongly expressed in differentiating neurons labelled for the early neuronal marker βIII tubulin (TUBB3, also known as Tuj1; a component of neuronal microtubules) throughout the dorso-ventral axis of the spinal neural tube (Fig. 1 B). This also revealed an uneven subcellular distribution of neuronal Gαi2, which was enriched in axons and sparse in apical processes (Fig. 1 B). To further characterise the subcellular distribution of Gαi2, TUBB3-positive neurons of the dorsal and medial spinal cord, which project their axons ventrally (*C*) were classified into stages according to morphology (Fig. 1 C). Neurons without axon outgrowth were classed as ‘pre-initiation’. Pre-initiation neurons with an apical process that extended to the ventricular surface were classified as ‘attached’, and they were classed as ‘delaminating’ if their apical process was partially or fully withdrawn. Neurons with a nascent axon (a ventrally oriented thick protrusion) shorter than approximately three cell body lengths were classified as ‘initiating’, while neurons with an axon longer than this were classed as ‘extending’. In attached neurons, Gαi2 expression was low and visually similar to the surrounding neuroepithelium (Fig. 1 D). In contrast, delaminating neurons exhibited a notable accumulation of Gαi2 at the baso-ventral side of the cell body (Fig. 1D). In initiating cells or cells extending axons Gαi2 accumulated in the nascent axon (Fig. 1 D, E) and was enriched in the growth cone (Fig. 1 E, F), where it was confined to the central zone (Fig. 1 G, H). Throughout differentiation, Gαi2 accumulations appeared set back from the border of the cell in relation to microtubules (Fig. 1 D, G, H), suggesting that Gαi2 accumulates intracellularly, away from the plasma membrane. These unexpected findings raised that possibility that polarised distribution of Gαi2 may play a role in facilitating axon initiation and outgrowth.

**Figure 1:**
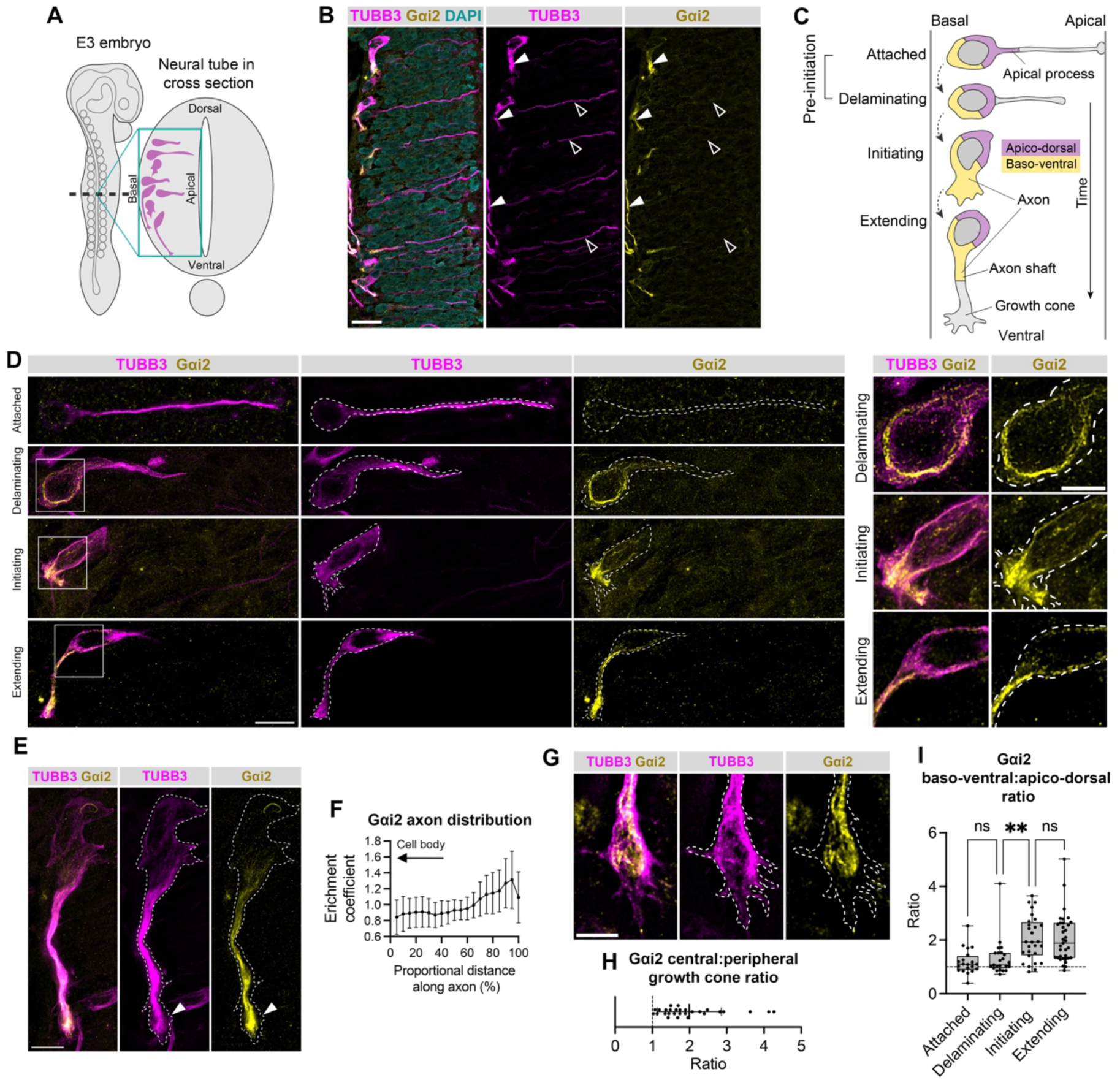
Gαi2 accumulates towards the fronts of differentiating neurons. **(A)** Schematic of E3 chick embryo showing differentiating neurons (purple) in a cross section of the spinal cord neural tube. **(B)** Immunofluorescence images of E3 chick trunk tissue showing Gαi2 distribution in the neural tube. Gai2 accumulates in differentiating neurons (indicated by TUBB3 labelling) in the basal tissue, is concentrated to axons (solid arrowheads), but is sparse in apical processes (empty arrowheads). Image representative of n=6 embryos, 101 cells. Note that some cells or axons are incomplete as a result of tissue sectioning. Scale bar=20 μm.**(C)** Schematic of morphological stages of spinal cord neuronal differentiation showing regions of interest used for analysis of Gαi2 distribution. **(D)** Immunofluorescence images of Gαi2 distribution in differentiating neurons. Neurons were staged according to their morphology using TUBB3 labelling. Gαi2 accumulation was low in attached neurons and, unlike evenly distributed TUBB3, accumulated towards the baso-ventral regions (fronts) of delaminating, initiating, and extending neurons. Gαi2 accumulations were set back from the border of the cell. Representative of attached n=20 cells, 6 embryos, delaminating n=23 cells, 9 embryos, initiating n=28 cells, 11 embryos, extending n=35 cells, 8 embryos. Left panel shows whole cells; scale bar=10 μm. Boxes indicate regions enlarged in right panel. Right panel scale bar=5 μm. **(E)** Immunofluorescence image showing axonal Gαi2 distribution. Gαi2 is found along the axon shaft and concentrated to the growth cone (arrowhead). Scale bar=10 μm. **(F)** Enrichment of Gαi2 along axon from proximal to distal in relation to the cell body (n=31 cells, 3 embryos). Graph shows means ± SD. **(G)** Immunofluorescence image showing Gαi2 accumulation in the central growth cone. **(H)** Quantification of Gαi2 fluorescence ratio between central and peripheral (0.5 μm-wide border) growth cone. N=28 cells, 9 embryos. **(I)** Gαi2 fluorescence intensity baso-ventral:apico-dorsal ratio. Baso-ventral region includes baso-ventral cell body and proximal axon or initiating process, where relevant. Apico-dorsal includes apico-dorsal cell body and apical process, where relevant. Attached n=20 cells from 6 embryos; delaminating n=23 cells from 9 embryos; initiating n=28 cells from 11 embryos; extending n=35 cells from 8 embryos. Kruskal-Wallis test with Dunn’s multiple comparisons, attached *vs* delaminating p>0.9999; delaminating *vs* initiating p=0.0011; initiating *vs* extending p>0.9999. Box plot in panel **I** shows median, range, and IQR. Other graphs show means ±SD. All images are maximum-intensity Z-projected stacks. **p≤0.01. White dashed lines indicate cell outlines.

To investigate if Gαi2 distribution was polarised along the axis of axon formation, regional Gαi2 fluorescence intensity was determined for cells at each stage of differentiation. Given that in the developing spinal cord, axons are associated with the baso-ventral cell body (Fig. 1 B, D, F), we partitioned the cell body into baso-ventral and apico-dorsal regions defined by tissue anatomy (Fig. 1 C) to approximate ‘front’ and ‘rear’ regions, respectively. The baso-ventral:dorso-apical ratio of Gαi2 fluorescence was greater than one for all stages (Fig. 1 I), confirming that Gαi2 distribution was polarised towards the cell front before and during axon formation. The ratio for cells initiating axons was significantly higher than delaminating neurons (Fig. 1 I), indicating enhanced polarisation of Gαi2 towards the front of the cell during initiation of axon outgrowth. Together, these results reveal that Gai2 accumulates intracellularly and asymmetrically towards the fronts of differentiating neurons, suggesting a role in facilitating axon outgrowth.

### Gαi2 distribution matches that of the polarised neurofilament network

To explore the mechanism of Gαi2 function in differentiating neurons, we examined the spatial relationship of Gαi2 to other cellular components. Phalloidin labelling revealed that Gαi2 accumulations did not coincide with filamentous actin (Fig. S 1 A, A’, A’’), which was concentrated at the cell periphery. Since Gα proteins are processed, sorted, and trafficked through common endomembrane pathways and can also participate in canonical signalling on internal membranes (*28, 29*), we compared the distribution of major endomembrane compartments to that of Gαi2. Immunolabelling of cells extending axons confirmed that Gαi2 accumulations did not coincide with the Golgi apparatus (labelled with anti-GM130 antibody; Fig. S1 B), or the endoplasmic reticulum (Fig. S1 C; labelled with anti-calreticulin antibody). Finally, early, late, and recycling endosomes (Fig. S1 D; labelled with anti-Rab5a, anti-Rab7a, and anti-CD71 antibodies respectively) were dispersed in the cell and not concentrated to the axon. These results suggest that neuronal intracellular Gαi2 is not in a state of processing, sorting, or trafficking, or participating in endomembrane signalling. Given that Gαi2 was set back from the border of the cell (Fig. 1 D, G, H) and not found to be associated with endomembranes (Fig. S1 B–D), our findings point towards a non-canonical mode of Gαi2 function in differentiating neurons.

Given that Gαi proteins have been shown to associate with and regulate microtubules (*21–25, 30*) and our finding that Gαi2 is concentrated within the microtubule-rich axon, we next examined the association between Gαi2 and microtubules. Following labelling for Gαi2 and TUBB3 (to identify neuronal microtubules), tissue was subjected to Airyscan super-resolution imaging. This revealed that Gαi2 accumulations had a filamentous form (Fig. 2 A, B), suggesting that Gαi2 might associate with a filamentous structure. In cells undergoing axon extension, longitudinal Gai2 accumulations aligned with microtubules in the axon shaft (Fig. 2 A), raising the possibility that Gαi2 associates with microtubules. However, the overall cellular distributions of Gαi2 and TUBB3 were different. Whereas Gαi2 was concentrated to the extending axon, TUBB3 was distributed more evenly throughout the cell (Fig. 2A,B). In addition, in some pre-initiation neurons, we observed distinct coiled Gai2 formations that clearly did not align with microtubules (Fig. 2 B). Together, these results indicated that Gai2 is associated closely with the microtubule network in the axon, but likely also associates with another filamentous structure in the cell.

**Figure 2:**
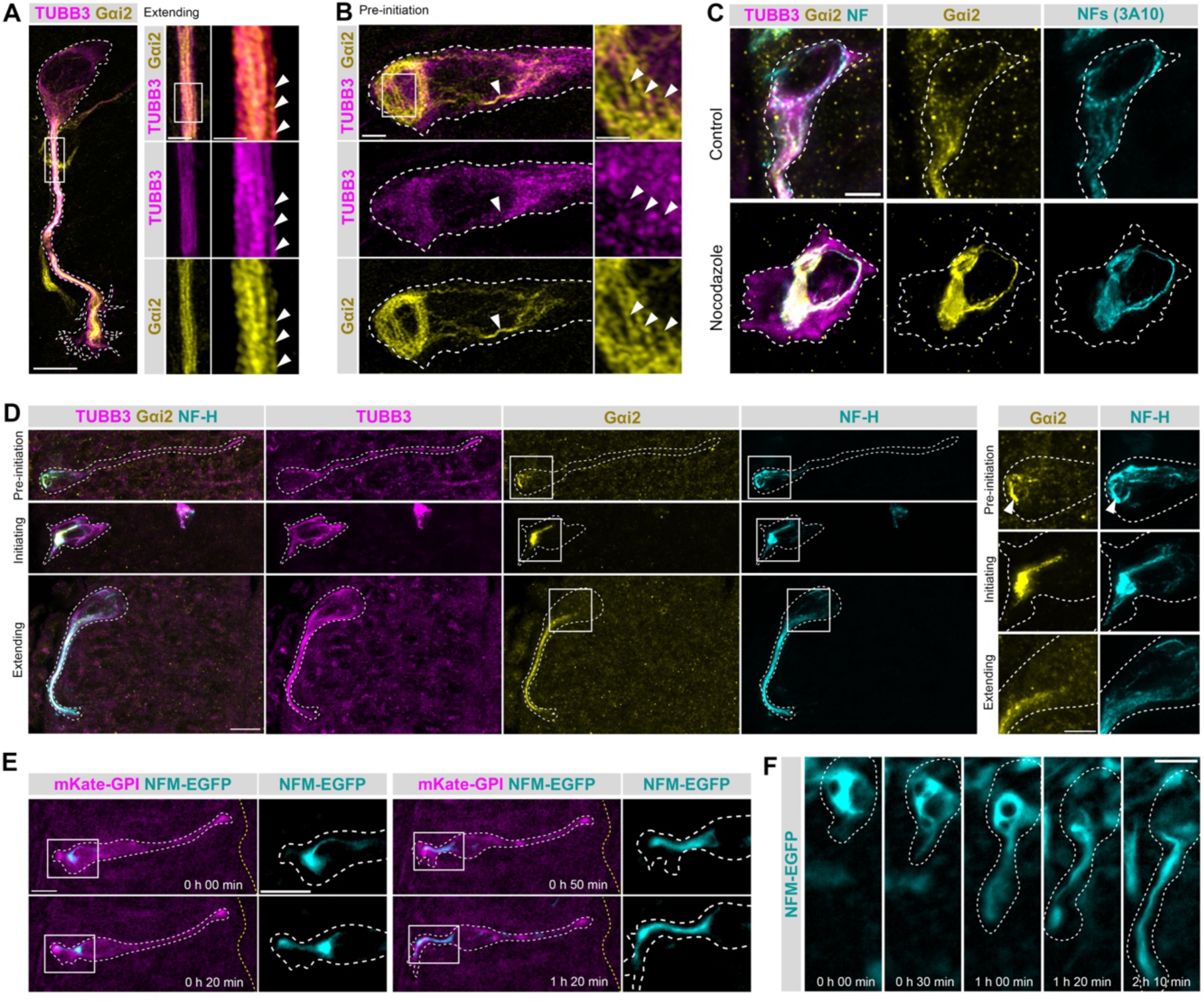
Gαi2 coincides with neurofilaments, which accumulate asymmetrically in differentiating neurons. **(A)** Airyscan immunofluorescence image of an extending neuron showing Gαi2 accumulating along microtubules (arrowheads), representative of n=49 cells from 5 embryos. Box on left image indicates region expanded in centre panel. Box on centre panel indicates region expanded in right panel. Scale bars: left=10 μm, centre=2 μm, right=1 μm. **(B)** Airyscan immunofluorescence image of a pre-initiation neuron showing coiled Gαi2 formations, not reflected in the tubulin architecture (TUBB3). Arrowheads indicate strand-like formations of Gαi2 not observed in TUBB3 labelling. Box shows expanded region in right hand panel. Scale bars from left to right 2 μm, 1 μm. **(C)** Immunofluorescence images of neurons in embryos treated for 2 h with high-dose (40 μM) nocodazole to depolymerise microtubules or DMSO alone (vehicle control). Gαi2 coincides with neurofilaments (labelled with 3A10 antibody) in both nocodazole-treated (n=101 cells, 3 embryos) and DMSO conditions (n=104 cells, 3 embryos). Scale bar=5 μm. **(D)** Immunofluorescence images of unmanipulated tissue showing distribution of Gαi2 and neurofilaments in differentiating neurons. Left panel shows whole neurons, staged by morphology. Neurofilaments are concentrated to the baso-ventral side of the cell (pre-initiation n=23/24 cells, 5 embryos; initiating 15/15 cells, 4 embryos; extending 52/53 cells, 5 embryos). Gαi2 accumulations are confined to regions enriched for neurofilaments (pre-initiation n=24 cells, 5 embryos; initiating 15 cells, 4 embryos; extending 53 cells, 5 embryos). White boxes indicate expanded regions in the right-hand panel. Arrowhead indicates a coiled formation of neurofilaments and Gαi2. Scale bars: left=10 μm, right=5 μm. **(E,F)** Live imaging showing neurofilament dynamics during axon formation in cells expressing NFM-EGFP. Real time from first frame shown indicated on each image. **(E)** Timelapse sequence of differentiating neuron expressing NFM-EGFP and mKate2-GPI undergoing axon formation showing neurofilament dynamics. Yellow dashed line approximates apical surface of tissue. Boxes indicate regions expanded in corresponding images to the right. As the axon emerges, the neurofilament accumulation extends into the protrusion (representative of n=33 cells, 6 embryos). Scale bars=10 μm. **(F)** Timelapse sequence of cell expressing NFM-EGFP showing unfurling of neurofilaments in a coiled formation which then extend into the axon (representative of n=27 cells, 6 embryos). Scale bar=10 μm. White dashed lines indicate cell outlines. All images are maximum-intensity Z-projected stacks.

The microtubule network controls the distribution of the many proteins, organelles, and cytoskeletal filaments that it associates with (*31*). To examine whether Gαi2 distribution was dependent on microtubules, we assessed Gαi2 distribution following microtubule depolymerisation by exposure of tissue to nocodazole for 2h. This resulted in an increased incidence of neurons without axons compared with tissue incubated in medium containing DMSO carrier control (Fig. S2). Furthermore, microtubule depolymerisation resulted in a high incidence of dense, coiled formations of Gαi2 in neuronal cell bodies compared with DMSO control (Fig. S2 A), which appeared similar to the coils we observed in unmanipulated delaminating neurons (Fig. 2B). These coil-like formations of Gαi2 were similar in shape to the collapsed intermediate filament networks that form after microtubule depolymerisation, as reported for vimentin or keratin in cultured epithelial cells or fibroblasts (*32–35*). Immunolabelling for neurofilaments (the dominant neuronal intermediate filament type) confirmed that Gαi2 accumulations coincide with neurofilaments (labelled with antibody 3A10), in nocodazole-treated and DMSO control cells (Fig. 2 C), suggesting that Gαi2 associates with the neurofilament network, and that the distribution of Gαi2 and neurofilaments is regulated by the microtubule network.

To investigate whether polarisation of Gαi2 accumulation in neurons was associated with polarised neurofilament accumulation, unmanipulated tissue was labelled for Gαi2, TUBB3, and the neurofilament protein NF-H. This confirmed that neurofilament distribution was polarised towards the baso-ventral side of the cell throughout differentiation (Fig. 2D). Neurofilaments were enriched in the axon and restricted to the central growth cone (Fig. S3), as previously reported (*3C*), and Gαi2 accumulations were confined to regions enriched for neurofilaments (Fig. 2D, S4). Additionally, this revealed that coils of Gαi2 were associated with neurofilament coils (Fig. 2D). These results reveal that neurofilaments form a polarised network in differentiating neurons and suggest that the polarised and filamentous distribution of Gαi2 occurs through association with the neurofilament network.

Given that little is known about the role of neurofilaments in axon initiation, we next tracked neurofilament behaviour in differentiating neurons using timelapse imaging. Neuroepithelial cells were transfected with the fluorescently tagged neurofilament protein construct EGFP-NFM (enhanced green fluorescent protein-NF-M) to label neurofilaments, mKate-GPI to label the cell membrane and Neurog2 to promote neuronal differentiation. The initial stages of neuronal differentiation were then monitored in *ex-vivo* slice cultures by timelapse imaging. We observed that as axon outgrowth initiated, dense neurofilament accumulations at the baso-ventral region extended into the core of the nascent axon (Fig. 2E; Movie 1). Where neurofilament coils were apparent, an unfurling motion was observed as the filaments extended (Fig. 2 F; Movie 2). Together, these results raised the possibility that neuronal Gαi2 is associated with microtubules and scaffolded by neurofilaments, which accumulate towards the fronts of differentiating neurons and unfurl into the initiating axon.

### Gαi2 associates with neurofilaments and microtubules

The microtubule and neurofilament networks are intimately connected (*37*). Given our observation that Gαi2 coincides with neurofilaments (Fig. 2 C, D) and accumulates along microtubules (Fig. 2 A), we hypothesised that Gαi2 may associate with both networks. To assess the association between Gαi2, neurofilaments, and microtubules, we performed *in-situ* proximity ligation assays (PLAs). In this method, conjugated primary antibody pairs bound to targets emit fluorescent signals when the targets are in close proximity to one another, enabling visualisation of protein–protein interactions (*38*). This revealed interactions between Gαi2 and NF-H, Gαi2 and TUBB3, and NF-H and TUBB3 in differentiating neurons, as indicated by PLA signal in the basal neuroepithelium and axons (Fig. S4). This is consistent with previous reports of interaction between microtubules and neurofilaments(*39-42*) and suggested that Gαi2 could interact with both networks.

To directly assess the arrangement of Gαi2, neurofilaments, and microtubules in neurons, we labelled tissue for Gαi2, NF-H, and TUBB3 and performed super-resolution lattice structured illumination microscopy (SIM). The proximal axon was examined, since this area is enriched for all three elements, but sparse enough to resolve filaments. This revealed a pattern of intertwining microtubules and neurofilaments in the nascent axon, decorated with Gαi2 (Fig. 3 A). Most Gαi2 was associated with neurofilaments and often overlapped with microtubules where neurofilaments were also present. Occasionally, we observed similar formations of Gαi2 not associated with either network, suggesting that Gαi2 might also associate with another structure. These results demonstrated that Gαi2 associates with the interconnected neurofilament and microtubule networks in differentiating neurons. Given that we found Gαi2 at points of overlap between both networks, we examined this relationship in more detail by generating a new image channel from colocalised NF-H and TUBB3 signal to isolate microtubule– neurofilament contact points. This revealed Gαi2 at and around colocalisation sites (Fig. 3 B), suggesting that Gαi2 is present at contact points between neurofilaments and microtubules.

**Figure 3:**
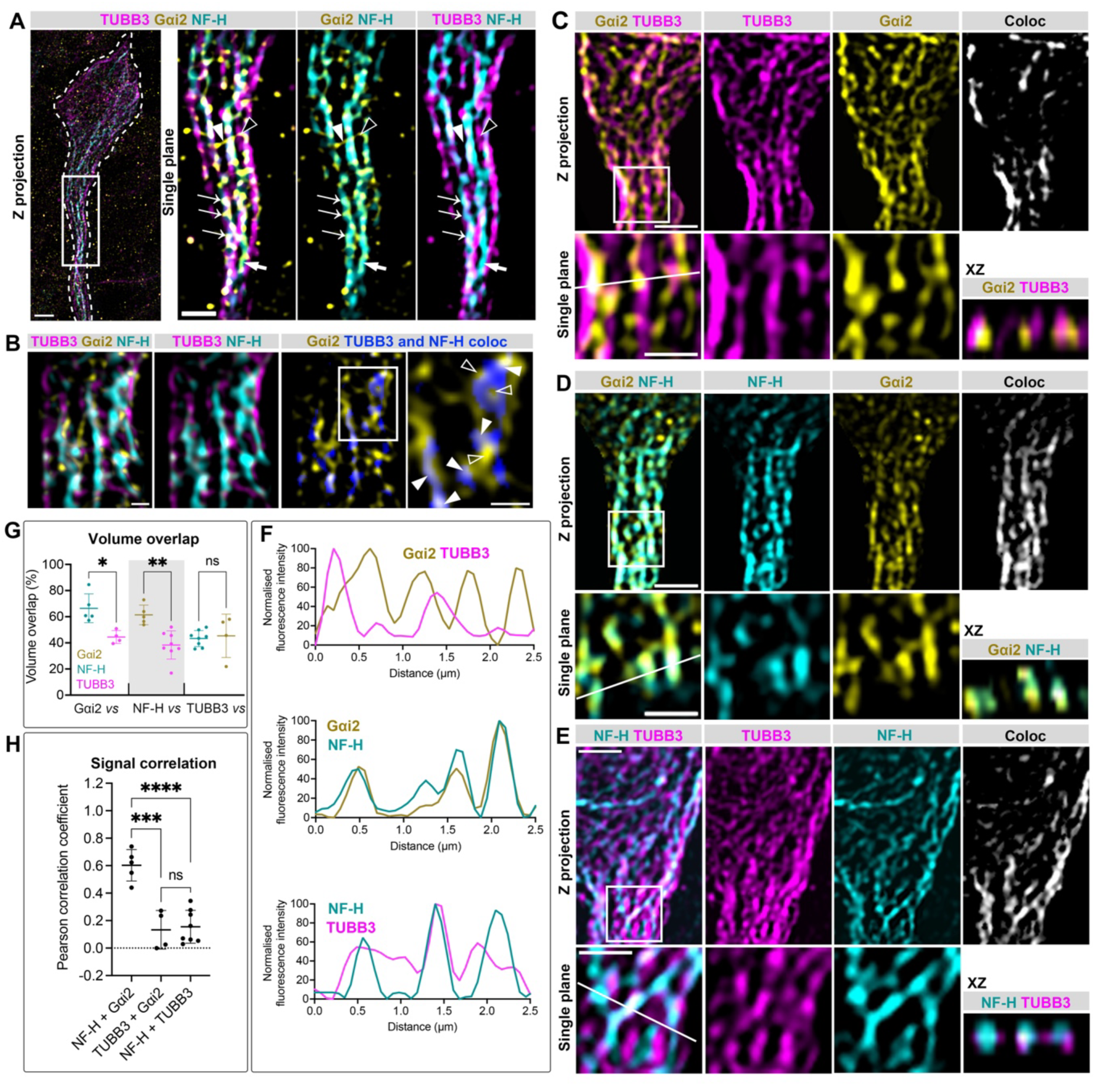
Gαi2 associates with the neurofilament and microtubule networks. Super-resolution microscopy immunofluorescence images showing association between Gαi2, neurofilaments (NF-H) and microtubules (TUBB3). **(A)** Lattice SIM images of an extending neuron showing Gαi2 decorating the intertwining neurofilament (NF-H) and microtubule (TUBB3) networks (thin arrows). Gαi2 is also present on neurofilaments alone (thick arrow), microtubules alone (black arrowhead), and not in association with either neurofilaments or microtubules (solid arrowhead). Left-hand image shows maximum-intensity projection. Box indicates region expanded as a single plane in left-hand panels. Scale bars: right=2 μm, left=1 μm. **(B)** Lattice SIM images of an axon shaft showing Gαi2 associating with contact points between microtubules and neurofilaments, as visualised by generation of a TUBB3 and NF-H colocalisation channel. Solid arrowheads indicate Gαi2 overlap with NFH–TUBB3 colocalisation. Box indicates region expanded in left-hand panel. Empty arrowheads indicate Gαi2 adjacent to NFH–TUBB3 colocalisation. Scale bars=500 nm. **(C–E)** STED microscopy images and analysis of colocalisation between Gαi2, neurofilaments, and microtubules in axons. In each panel, top row shows maximum-intensity Z-projection of proximal axon; far-right panel shows colocalisation channel generated from overlap between the two channels; box indicates region expanded below as a single plane; white line indicates region of interest for corresponding profile plot in panel F; bottom right section in orthogonal (XZ) view. Scale bars: top panel=2 μm, expanded region=1 μm. **(C)** Gαi2 and microtubules. **(D**) Gαi2 and neurofilaments. **(E)** Neurofilaments and microtubules. **(F)** Fluorescence intensity profiles for pairs of targets taken from linear region of interest specified in bottom right images of **C–E**. **(G)** Percentage overlap between 3D volumes in STED images. Gαi2 with NF-H n=5 cells from 3 embryos. Gαi2 with TUBB3 n=4 cells from 2 embryos. NF-H with Gαi2 n=5 cells from 3 embryos. NF-H with TUBB3 n=8 cells from 3 embryos. TUBB3 with Gαi2 n=4 cells from 2 embryos. TUBB3 with NF-H n=8 cells from 3 embryos. One-way ANOVA with Tukey’s multiple comparisons test, Gαi2 with NFH vs Gαi2 with TUBB3 p=0.0250; NF-H with Gαi2 vs NF-H with TUBB3 p=0.9993; TUBB3 with NF-H vs TUBB3 with Gαi2 p=0.9993. **(H)** Correlation of signal covariance within colocalised regions expressed as Pearson’s correlation coefficient. NF-H and Gαi2 n=5 cells from 3 embryos. TUBB3 and Gαi2 n=4 cells from 2 embryos. NF-H and TUBB3 n=8 cells from 3 embryos. One-way ANOVA with Tukey’s multiple comparisons test, NFH and TUBB3 vs TUBB3 and Gαi2 p=0.9546; NF-H and TUBB3 vs NF-H and Gαi2 p<0.0001; TUBB3 and Gαi2 vs NF-H and Gαi2 p=0.0002. White dashed lines indicate cell outlines. Graphs **G** and **H** show means ± SD. *p≤0.05. **p≤0.01. ***p≤0.001. ****p≤0.0001.

To compare the degree of association between components, we conducted STED super-resolution imaging of fixed tissue labelled for pairs of targets and performed colocalisation analysis. This confirmed colocalisation of signal between all pairs of components (Fig. 3 C–E). Fluorescence intensity profiles of Gαi2 and NF-H signal along a linear region of interest (indicated in Fig. 3 C–E) revealed matching peaks of fluorescence intensity between Gαi2 and NF-H, whereas plots of Gαi2 with TUBB3 and NF-H with TUBB3 exhibited overlapping but more offset peaks (Fig. 3 F). Additionally, correlation of signal covariance and volume overlap between Gαi2 and NF-H was significantly higher than between Gαi2 and TUBB3 or NF-H and TUBB3, which overlapped to a similar degree (Fig. 3 G, H). Together, these results are consistent with Gαi2 accumulating on neurofilaments and associating at points of contact with microtubules, suggesting a role for Gαi2 in coordinating these networks during neuronal differentiation.

### The microtubule network is polarised towards the site of axon outgrowth

Differentiating spinal cord neurons reorient their polarity axis away from the apico-basal polarity of their precursor cells to establish an axon ventrally (*C*). We found that this polarisation event is associated with asymmetric accumulation of Gαi2-decorated neurofilaments towards the axon (Figs 1,2). However, the mechanisms governing polarity in de-novo axon formation are unclear. The organisation of the microtubule network directs cell polarity by distributing cellular components and controlling cell shape (*8*). Network asymmetry orients polarity and can be achieved through positioning of microtubule-organising centres (MTOCs), such as the centrosome. To investigate whether the microtubule network instructs polarity in differentiating neurons, we monitored the positioning of the centrosome during early neuronal differentiation. Neuroepithelial cells were transfected with PACT-TagRFP to label the centrosome, GFP-GPI, and Neurog2, and the resulting differentiating neurons were monitored through timelapse imaging. Consistent with previous observations (*5, 43*), the centrosome was located at the tip of the retracting apical process during delamination (Fig. 4 A; Movie 3). Upon completion of apical process retraction, the centrosome was delivered to the cell body and was positioned opposite the site of the axon at the moment of initiation (Fig. 4 A, B; Movie 3) before moving towards the base of the extending axon (Movie 3). To confirm the orientation of microtubules, we transfected neuroepithelial cells with EB3-GFP to label microtubule plus tips, dsRed-Cent2 to label the centrosome, and Neurog2, and conducted high-speed timelapse imaging. This facilitated visualisation of microtubule growth trajectories, which were observed to emanate from the region of the centrosome towards the nascent axon (Fig. 4 C; Movie 4). These results suggested that the centrosome acts as an MTOC in early differentiating neurons and demonstrate that microtubule network orientation correlates with the axis of axon formation.

**Figure 4:**
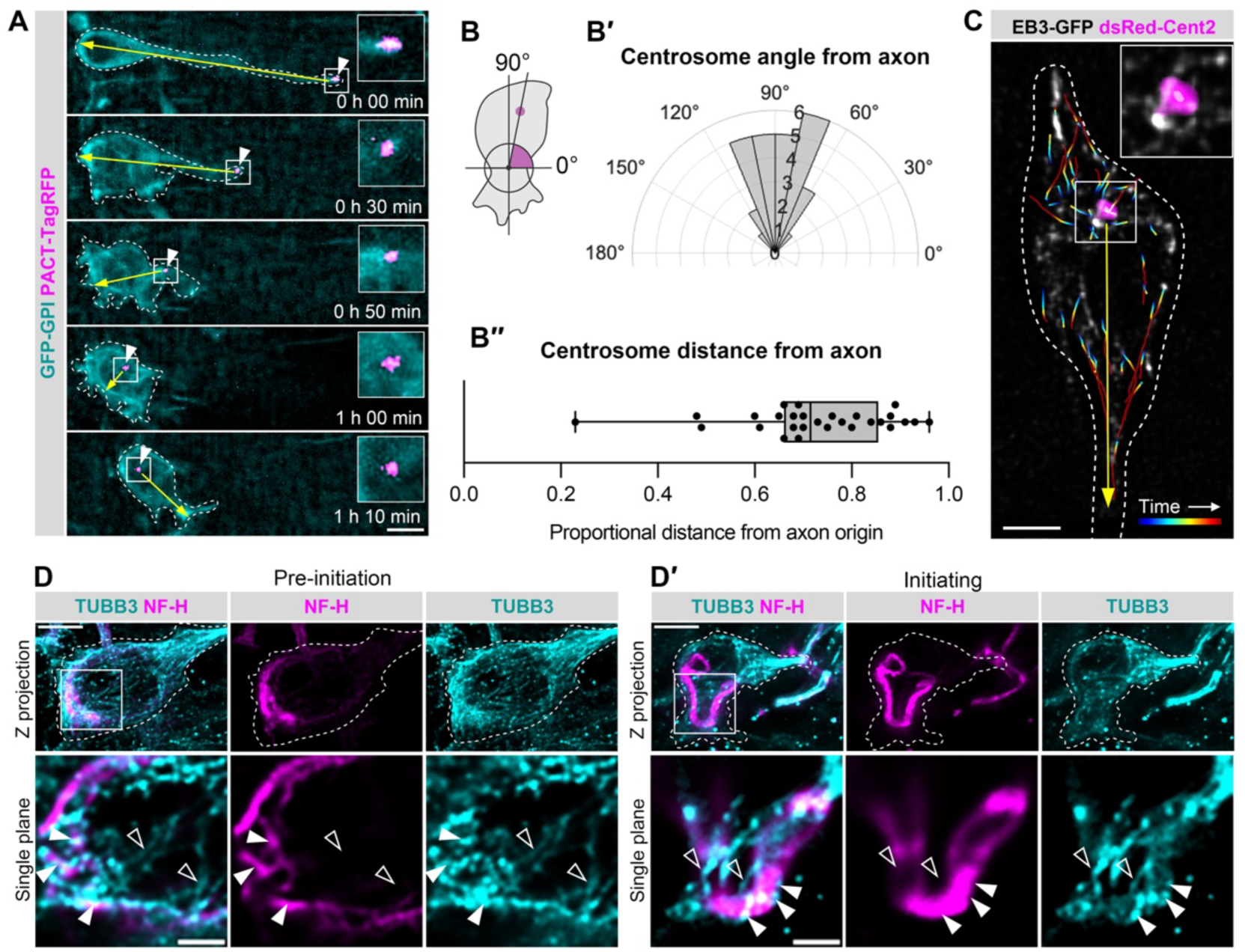
The microtubule network orients towards the site of axon formation. **(A)** Timelapse sequence of differentiating neuron expressing PACT-TagRFP and GFP-GPI showing centrosome positioning before and during axon formation. Arrowheads indicate centrosome. Boxes indicate expanded regions shown in insets. In the delaminating stage, the centrosome is in the tip of the apical process (n=11 cells, 9 embryos). Yellow arrow traces axis from centrosome through the cell body centre to the opposite side of the cell. Note that the axis shifts as the centrosome enters the cell body and an axon is initiated. Images acquired every 10 min. Scale bar=10 μm. **(B–B′′)** Quantification of centrosome position at axon initiation from timelapse imaging. **(B)** Schematic showing measurement of angle from axon initiation site, with 0° perpendicular and apical to the axon and 90° directly opposite. Purple dot indicates centrosome. **(B′)** Polar histogram of centrosome angle at the point of axon initiation (n=28 cells, 15 embryos). **(B′′)** Distance of centrosome from axon origin site shown as a proportion of total length of cell body. Median distance 0.72 (IQR 0.66–0.86). Graph shows median, range, and IQR. **(C)** Approximation of microtubule network orientation from microtubule growth trajectories shows microtubules emerge from the region of the centrosome and track towards and into the axon (representative of n=17 cells, 15 embryos). Frame from timelapse sequence of extending cell expressing EB3-GFP and dsRed-Cent2 showing centrosome position and microtubule plus-tip tracking. Images were acquired every 1.66 s. EB3-GFP comet trajectories shown in time-coded tracks on image. Boxed region expanded in inset shows centrosome. Scale bar=5 μm. **(D–D′)** Immunofluorescence images of differentiating neurons showing microtubules (TUBB3) associating with neurofilaments (NF-H) at the baso-ventral cell body of preinitiating (**D**; n=17 cells, 3 embryos) and initiating **(D**; n=7 cells, 3 embryos). Upper panels: maximum-intensity Z-projected whole-cell images. Boxes indicate zoomed regions in lower panels. Lower panels: single focal plane expanded region. Empty arrowheads indicate microtubules directed to the cell front. Solid arrowheads show microtubules contacting neurofilaments. Scale bars: upper=5 μm, lower expanded images=2 μm. White dashed lines indicate cell outlines.

Microtubules control intermediate filament network distribution, including that of neurofilaments (*44-46*). We found that neurofilaments accumulate in coiled formations in the cell body and unfurl into the axon as it emerges (Fig. 2 D–F). However, the mechanism by which neurofilaments transition into the initial axonal protrusion is unclear. To explore the relationship between microtubules and neurofilaments as neurofilaments begin to enter the nascent axon, we labelled fixed tissue for TUBB3 and NF-H. In pre-initiation cells, we observed microtubules (linear formations of TUBB3) emerging from the apico-dorsal region of cells (eg, the apical process) and overlapping with neurofilaments at the baso-ventral region (Fig. 4 D). In cells initiating axons, microtubules were observed to enter the protrusion accompanied by extended neurofilament accumulations, which were set back from microtubules at the periphery (Fig. 4 D′) raising the possibility that microtubules draw neurofilaments into the nascent axon. Together, these findings suggest that the microtubule network directs polarisation of cellular structure during axon initiation. Furthermore, given our finding that Gαi2 associates with points of contact between microtubules and neurofilaments, these observations raise the possibility that Gαi2 may coordinate microtubule-driven entry of neurofilaments into the initiating axon.

### Gαi2 regulates passage of neurofilaments into initiating axons

To investigate the role of Gαi2 in differentiating neurons, Gαi2 protein levels were depleted by performing RNA interference (RNAi)-mediated knockdown (*47*). Neural progenitor cells were transfected with a construct expressing short hairpin RNA targeting Gαi2, and RFP as a marker for transfection (pRFPRNAi-Gαi2) or a control construct targeting firefly luciferase (pRFPRNAi-Luc). Following incubation for 26h, embryos were fixed and labelled for TUBB3 and Gαi2 to assess knockdown efficiency. pRFPRNAi-Gαi2 expression resulted in a significant reduction of axonal Gαi2 normalised mean fluorescence intensity when compared with cells expressing pRFPRNAi-Luc (Fig. S5). Interestingly, we noted that knockdown of Gαi2 led to an increase in the proportion of pre-initiation and initiating cells relative to extending cells (Fig. 5 A, B), suggesting that Gαi2 depletion disrupted stable axon formation. This led us to investigate the mechanistic basis through which this may be taking place.

**Figure 5:**
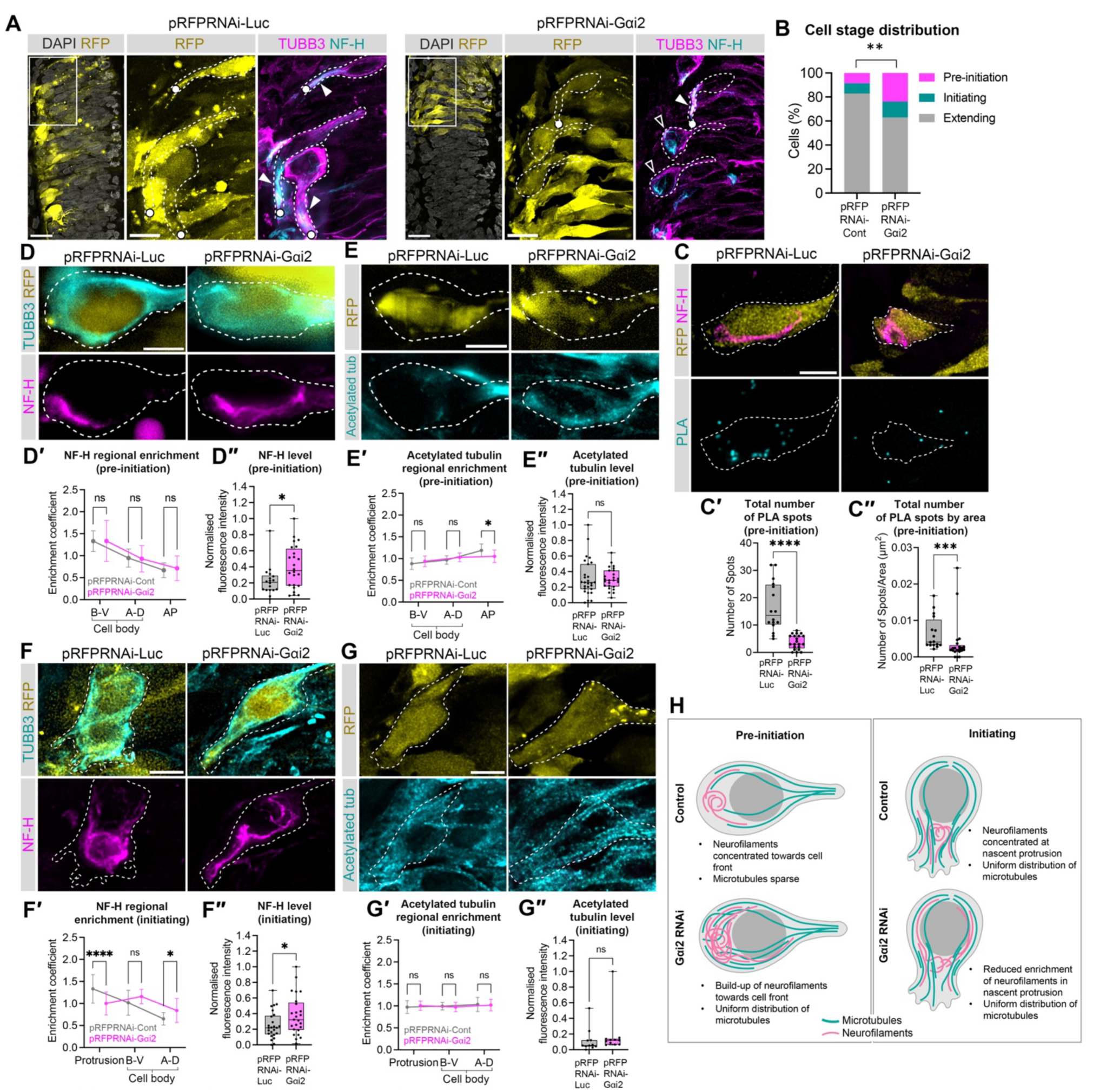
Gαi2 regulates neurofilament distribution in differentiating neurons. **(A)** Immunofluorescence images showing stages of differentiating neurons expressing pRFPRNAi-Luc or pRFPRNAi-Gαi2. Transfected cells indicated by cytoplasmic RFP expression. Left-hand images show one side of the neural tube tissue in cross-section. Scale bars=20 μm. Box indicates region expanded in centre and right-hand images. Expanded images scale bars=10 μm. Arrowheads indicate axons. Circles indicate axons incomplete due to tissue sectioning. **(B)** Proportion of transfected cells at defined morphological stages in tissue electroporated with pRFPRNAi-Luc or pRFPRNAi-Gαi2. Fisher’s exact test p=0.0033 (pRFPRNAi-Luc n=106 cells, 3 embryos; pRFPRNAi-Gαi2 n=105 cells, 4 embryos). **(C)** Immunofluorescence images of *in-situ* proximity ligation assays for interactions between neurofilaments (NF-H) and microtubules (TUBB3) in cells expressing pRFPRNAi-Luc or pRFPRNAi-Gαi2 and labelled for NF-H. **(C’)** Number of PLA spots counted at the cell front of pre-initiating neurons. Mann-Whitney test: p<0.0001 (pRFPRNAi-Luc n=16 cells, 4 embryos; pRFPRNAi-Gαi2 n=21 cells, 4 embryos). **(C’’)** Number of PLA spots counted at the cell front of pre-initiating neurons by area. Mann-Whitney test: p=0.0008 (pRFPRNAi-Luc n=16 cells, 4 embryos; pRFPRNAi-Gαi2 n=21 cells, 4 embryos). **(D-G)** Neurofilaments and microtubule network distributions in neurons expressing pRFPRNAi-Luc or pRFPRNAi-Gαi2. Transfected cells indicated by cytoplasmic RFP expression. **(D-D′′)** Neurofilament distribution in pre-initiation cells**. (D)** Immunofluorescence images of pre-initiation cells labelled for neurofilaments (NF-H) and TUBB3 (neuronal microtubules/cell shape). **(D′)** Enrichment of neurofilaments in regions of interest. Mixed-effects model with Geisser-Greenhouse correction and Šidák’s multiple comparisons test: baso-ventral cell body p>0.9999; apico-dorsal cell body p=0.9956; apical process p=0.9066 (pRFPRNAi-Luc cell body n=18 cells, 6 embryos, apical process n=15 cells, 5 embryos; pRFPRNAi-Gαi2 cell body n=24 cells, 4 embryos, apical process n=22 cells, 4 embryos). **(D′′)** Mean neurofilament levels across all regions of interest. Mann-Whitney test p=0.0268 (pRFPRNAi-Luc cell body n=18 cells, 6 embryos, apical process n=15 cells, 5 embryos; pRFPRNAi-Gαi2 cell body n=24 cells, 4 embryos, apical process n=22 cells, 4 embryos). **(E-E′′)** Microtubule distribution in pre-initiation cells. **(E)** Immunofluorescence images of pre-initiation cells labelled for stable microtubules (acetylated tubulin, Lys40). (**E′)** Enrichment of stable microtubules in regions of interest. Mixed-effects model with Geisser-Greenhouse correction and Šidák’s multiple comparisons test: baso-ventral cell body p=0.3442; apico-dorsal cell body p=0.0978; apical process p=0.0185 (pRFPRNAi-Luc cell body n=27 cells, 4 embryos, apical process n=22 cells, 3 embryos; pRFPRNAi-Gαi2 cell body n=25 cells, 3 embryos, apical process n=18 cells, 2 embryos). **(E′′)** Mean stable microtubule levels across all regions of interest. Mann-Whitney test p=0.5609 (pRFPRNAi-Luc cell body n=27 cells, 4 embryos, apical process n=22 cells, 3 embryos; pRFPRNAi-Gαi2 cell body n=25 cells, 3 embryos, apical process n=18 cells, 2 embryos). **(F-F′′)** Neurofilament distribution in initiating cells. **(F)** Immunofluorescence images of initiating cells labelled for neurofilaments (NF-H) and TUBB3 (neuronal microtubules/cell shape). **(F′)** Enrichment of neurofilaments in regions of interest. Two-way ANOVA with Šidák’s multiple comparisons: protrusion p<0.0008; baso-ventral cell body p=0.1082; apico-dorsal cell body p=0.0117 (each condition: n=25 cells, 5 embryos). **(F′′)** Mean neurofilament levels across all regions of interest. Unpaired t test p=0.0456 (each condition: n=25 cells, 5 embryos). **(G-G′′)** Microtubule distribution in initiating cells. **(G)** Immunofluorescence images of initiating cells labelled for stable microtubules (acetylated tubulin, Lys40). **(G′)** Enrichment of stable microtubules in regions of interest. Two-way ANOVA with Šidák’s multiple comparisons: protrusion p=0.9161; baso-ventral cell body p=0.9588; apico-dorsal cell body p=0.9963 (each condition: n=11 cells, 3 embryos). **(G′′)** Mean stable microtubule levels across all regions of interest. Mann-Whitney test p=0.0652 (each condition: n=11 cells, 3 embryos). **(H)** Schematic summarising the effect of Gαi2 depletion on neurofilament and microtubule distribution. White dashed lines indicate cell outlines. All images are maximum-intensity Z-projected stacks. Scale bars=5 μm. Graphs **D′, E′, F′** and **G′** show means ± SD. Graphs **C′, C′′, D′′, E′′, F′′** and **G’’** are visualised with median, IQR, and range to allow consistent visualisation across datasets. Statistical testing used parametric or nonparametric methods as appropriate (see Materials and methods). *p≤0.05. ****p≤0.0001. A-D=apico-dorsal. B-V=baso-ventral. AP=apical process.

Gαi2 was recently reported to regulate microtubule dynamics in migrating neural crest cells (*22*). Given the close association of neuronal Gαi2 with the microtubule network, and the importance of microtubules in neurite initiation (*2, 3, 48*), we speculated that Gαi2 might also modulate neuronal microtubule dynamics. However, depletion of Gαi2 did not affect the speed or distance travelled by EB3 comets in axons (Fig. S6), implying that Gαi2 does not affect microtubule growth dynamics in differentiating neurons.

As Gαi2 is present at points of microtubule–neurofilament contact (Fig. 3B) and microtubules associate with neurofilaments in early axon formation (Fig. 4 D), we hypothesised that Gαi2 may facilitate coupling of the neurofilament and microtubule networks to facilitate microtubule-mediated passage of neurofilaments into the initiating axon. To investigate this, we first assessed the effect of Gαi2 depletion on the interaction between microtubules and neurofilaments by performing *in-situ* proximity ligation assays. This technique revealed a drop in interactions between microtubules and neurofilaments in pre-initiation cells relative to control cells expressing the luciferase targeting construct (Fig. 5 C, C’, C’’), indicating reduced engagement between these networks. Next, we assessed the effect this had on microtubule and neurofilament network distribution in immunolabelled tissue. Given that a substantial amount of TUBB3 in neurons is not polymerised (*4S*), tissue was labelled for acetylated tubulin to identify tubulin stably incorporated into microtubules (*50*). Neurofilaments were identified by NF-H labelling. To determine distribution of the networks before and during axon formation, fluorescence intensity was measured in key cellular regions of pre-initiation and initiating neurons. Polarisation of distribution was then assessed by variation of regional enrichment. Consistent with our previous observations in unmanipulated tissue (Fig. 2D), in pre-initiation cells expressing the control RNAi construct, neurofilaments were concentrated towards the baso-ventral cell front (Fig. 5 D, D’). Depletion of Gαi2 resulted in increased overall levels of neurofilaments in the cell body, suggestive of a build-up of neurofilaments, but these remained asymmetrically distributed towards the cell front (Fig. 5 D, D’, D’’). Additionally, whereas microtubules were sparse towards the fronts of control pre-initiation cells, Gαi2 depletion resulted in a uniform distribution of microtubules in cells (Fig. 5 E, E’, E’’). These results suggested that Gαi2 depletion leads to a build-up of microtubules and neurofilaments at the cell front in pre-initiation cells. To investigate this further, we examined cells that had initiated axons. In control cells undergoing axon initiation, neurofilaments were concentrated at the nascent protrusion (Fig. 5 F, F’), whereas microtubules were evenly dispersed across the cell (Fig. 5 G, G’). Depletion of Gαi2 reduced enrichment of neurofilaments in the nascent protrusion and increased their enrichment at the apico-dorsal cell body (Fig. 5 F, F’), suggesting that neurofilament accumulation in the nascent protrusion was perturbed. Furthermore, overall levels of NF-H were increased (Fig. 5 F, F’, F’’), suggesting a build-up of neurofilaments when Gαi2 is depleted. Gαi2 depletion did not affect the distribution or levels of stable microtubules in initiating cells (Fig. 5 G, G’, G’’). These observations suggest that in Gαi2-depleted cells attempting to initiate an axon, microtubules invade the nascent protrusion, as in control cells, but the entry of neurofilaments is disrupted (Fig. 5 H). Together with our finding that microtubule depolymerisation increases the incidence of Gαi2-associated neurofilament coils in the neuronal cell body (Fig. 2 C), these results suggest that Gαi2 regulates microtubule-driven passage of neurofilaments into the axon by facilitating engagement of the microtubule and neurofilament networks. Furthermore, together with our observation that fewer Gαi2-depleted neurons have axons (Fig. 5 A, B), these results introduce the possibility that regulated transit of neurofilaments into axons is a prerequisite for axon formation.

### Gαi2 stabilises axon extension

Given that neurofilaments stabilise neurite outgrowth (*9*), we reasoned that perturbed neurofilament passage into nascent axons of Gαi2 depleted cells could result in protrusion instability and disrupted axon formation. To examine the effect of Gαi2 depletion on axon formation in real time, neuroepithelial cells were co-transfected with RNAi constructs, GFP-GPI, and Neurog2 and subjected to timelapse imaging. Whereas in the control condition, typically cells extended a single stable axon, Gαi2 depletion was associated with formation of multiple unstable protrusions (Fig. 6 A–C; Movie 5). In a 6.5-h period, the majority (75%) of control cells initiated one major protrusion; however, Gαi2 depletion resulted in the majority (67%) of cells initiating two or more (Fig. 6 A, B). When individual protrusions were tracked for 6.5h following initiation, Gai2 depletion significantly reduced the mean per-cell persistence of protrusions (Fig. 6 A, C). These results demonstrate that Gαi2 assists in the formation of a single stable axon. Together with our findings that Gαi2 associates with the neurofilament and microtubule networks and regulates the entry of neurofilaments into the nascent axon, our results suggest that by supporting microtubule-mediated transit of neurofilaments, Gαi2 facilitates formation of a stable axon.

**Figure 6:**
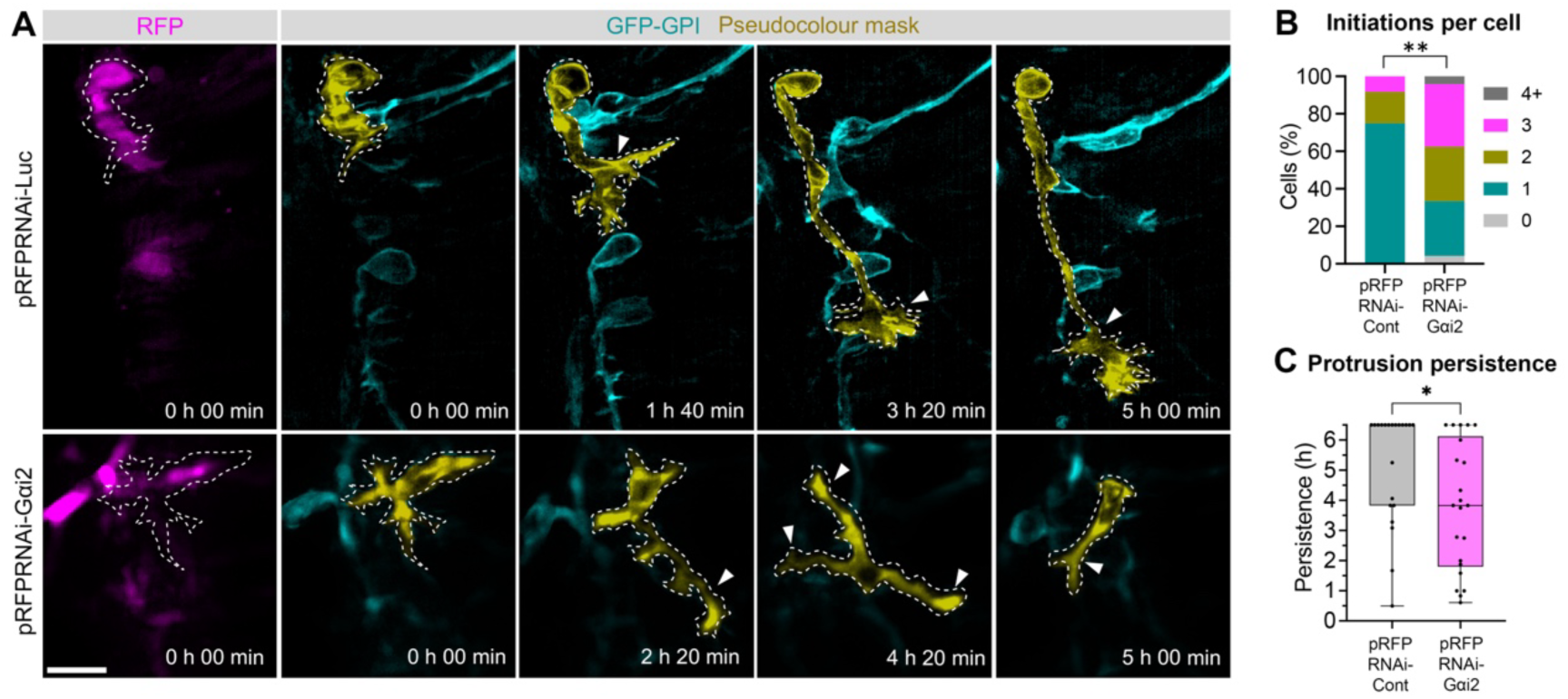
Gαi2 depletion disrupts axon outgrowth. **(A)** Timelapse sequence showing formation of major protrusions in differentiating neurons expressing pRFPRNAi-Luc or pRFPRNAi-Gαi2 and GFP-GPI. Cells expressing RNAi constructs are indicated by detection of cytoplasmic RFP. Images were acquired every 10 min. Real time from first image shown indicated for each image. Cell of interest is pseudocoloured to highlight its shape. Arrowheads indicate major protrusions. Scale bar=20 μm. **(B)** Number of protrusions initiated per cell within 6.5 h of tracking. Fisher’s exact test p=0.0099 (pRFPRNAi-Luc n=24 cells, 12 embryos; pRFPRNAi-Gαi2, n=24 cells,14 embryos). **(C)** Persistence of major protrusions tracked for 6.5 h from initiation. Mann-Whitney test p=0.0158 (pRFPRNAi-Luc n=21 cells, 12 embryos; pRFPRNAi-Gαi2, n=22 cells,14 embryos). White dashed lines indicate cell outlines. All images are maximum-intensity Z-projected stacks. *p≤0.05. **p≤0.01.

## Discussion

In this study, we demonstrate that *de-novo* axon formation *in vivo* is characterised by pre-polarisation of the cytoskeleton and that establishment of cytoskeletal and morphological polarity is locally regulated by the G protein Gαi2. We reveal that axon outgrowth is characterised by asymmetric organisation of the centrosome-derived microtubule network and asymmetric accumulation of Gαi2-associated neurofilaments towards the cell front. Here, microtubules contact neurofilaments, which then enter the nascent axon behind microtubules as it emerges. Importantly, Gαi2 accumulates on the neurofilament network and facilitates neurofilament association with intermingling microtubules. Gαi2 facilitates the accumulation of neurofilaments in the axon and ensures that axon formation is focused and persistent. Thus, we reveal previously unknown cytoskeletal dynamics during axon initiation and find a novel role for Gαi2 in establishment of cellular polarity and coordination of cytoskeletal networks.

Based on the data presented here, we propose the following model for cytoskeletal control of polarity and axon formation in spinal cord neurons (Fig. 7). In new-born neurons undergoing delamination from the neuroepithelium, the centrosome is located at the tip of the retracting apical process and directs microtubules towards the basal pole of the cell (Fig. 4 A–C). Consistent with previous observations that centrosome positioning can direct front–rear polarity (*51, 52*), delivery of the centrosome to the cell body may facilitate a basal-to-ventral shift in microtubule-network orientation. Concurrently, Gαi2-associated neurofilaments accumulate at the nascent baso-ventral cell front. Given that neurofilaments are subject to microtubule-based trafficking (*53*) and considering our finding that neurofilament distribution is reliant on microtubules (Fig. 2 C), this could be achieved by directed neurofilament transport towards microtubule plus-ends directed to the cell front. The axon is then initiated, likely initially through actin-dependent deformation of the cell membrane before microtubules enter the protrusion (*2, 3*). Since Gαi2 is present at points of microtubule–neurofilament contact (Fig. 3 B) and Gαi2 depletion results in both reduced interactions between microtubules and neurofilaments (Fig. 5 C) and disrupted axonal neurofilament accumulation (Fig. 5 F), we propose that Gαi2 facilitates engagement of microtubules with neurofilaments, resulting in accumulated neurofilaments at the cell front being drawn into the axon, stabilising the growing protrusion. Thus, depletion of Gαi2 results in a reduction in engagement of neurofilaments with microtubules, resulting in fewer neurofilaments passing into the protrusion, leading to the observed instability and retraction (Fig. 6). Reduced neurofilament passage into the protrusion plus retraction of unstable protrusions could result in the observed build-up of neurofilaments and microtubules at the cell body in Gαi2-depleted neurons (Fig. 5 D–F). Repeated attempts to form an axon could then manifest as the observed initiation of multiple unstable protrusions (Fig. 6).

**Figure 7:**
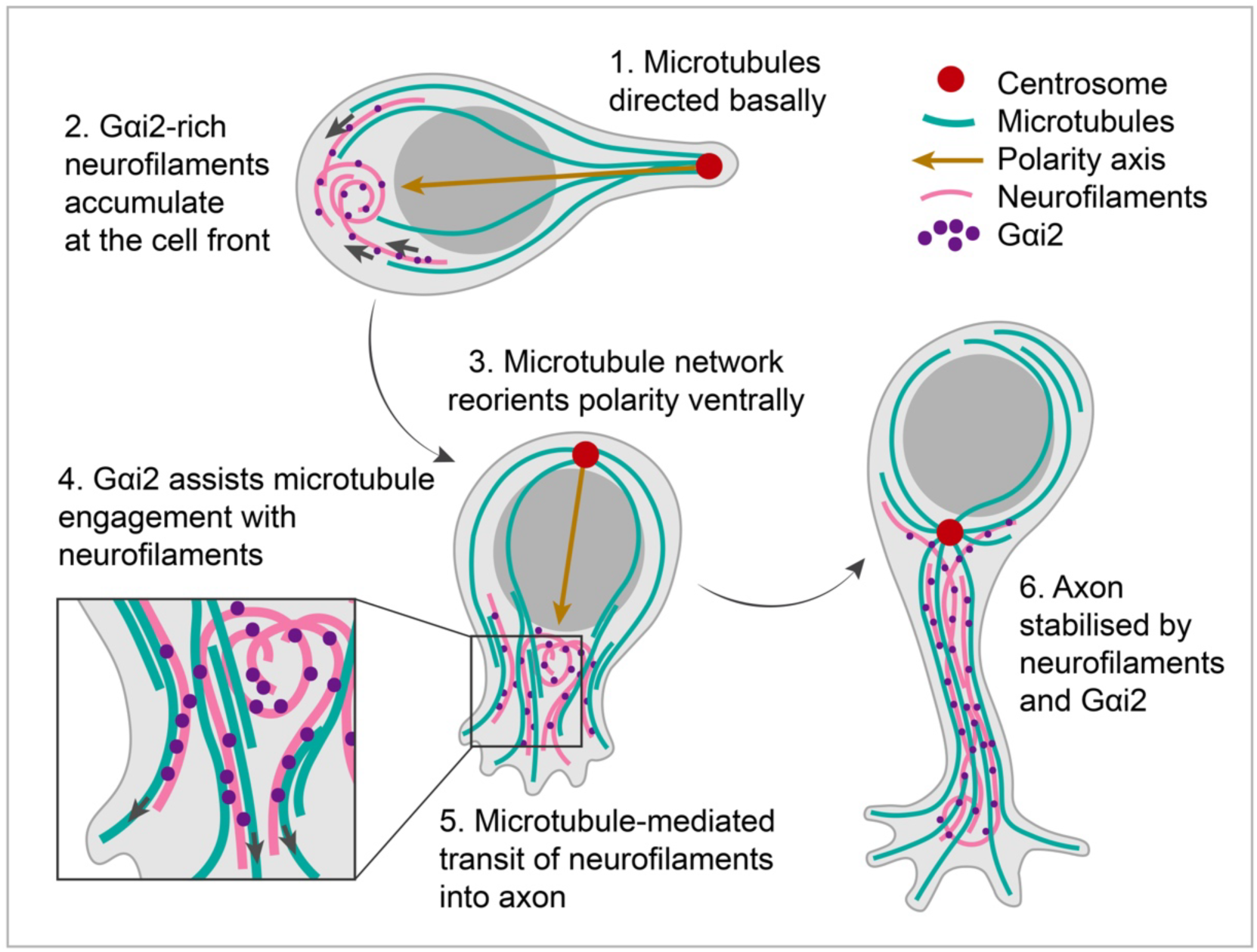
Model for cytoskeletal control of polarity and axon formation in spinal cord neurons.

Neurofilaments are known as the structural stabilisers of the axon (*46*); however, little is known about their role in axon initiation. Previous studies of cultured cells have suggested neurofilaments support neurite outgrowth, as demonstrated by perturbed growth and stability following disruption of neurofilaments (*54–57*). However, to our knowledge, this is the first study to examine neurofilament behaviour during axon initiation *in vivo*. We show that neurofilaments accumulate asymmetrically before axon initiation before populating the nascent axon as it is initiated (Fig. 2 D–F), suggesting that neurofilaments have a role in the establishment of neuronal polarity. This adds to previous work demonstrating that other intermediate filaments, such as vimentin, form asymmetric networks and aid polarity establishment in other cell types undergoing morphological transitions (*44, 58-60*). Due to their structural resilience (*9*), neurofilaments could maintain polarised changes to neuronal structure set up by microtubules, while also permitting remodelling of microtubules, as has been reported for the intermediate filament vimentin in migrating cells (*44*). We found that before axon formation, neurofilaments accumulate asymmetrically in the baso-ventral cell body (Fig. 2 D–F). Whereas microtubules are highly dynamic polymers (*61*), neurofilaments are stable and long-lived (*46*). Their gradual deposit at the cell front could favour axon formation from that region, for example by providing local structural reinforcement during microtubule and actin remodelling. Interestingly, the coiling of neurofilaments, which we observed in pre-initiation neurons (Fig. 2 D, F), has previously been observed in more mature neurons at axon branch points (*62*), where neurofilaments undergo stalling of movement, suggesting these coiled aggregates might form as they meet resistance before axon protrusion. Since neurofilaments display slow turnover (*46*), their aggregation at the cell front could serve as a store for immediate use in the new axon. Neurofilaments tethered to microtubules could be drawn from this deposit into the axon as it is populated by microtubules. Indeed, our demonstration that coiled neurofilaments unfurl into the axon (Fig. 2 F), much like twine unravelling from a ball, supports this idea.

Positioning of the centrosome directs the polarity of the microtubule cytoskeleton and therefore cellular organisation (*51, 52*). We demonstrate that the centrosome is located opposite the site of axon emergence in chick spinal cord neurons *in vivo* (Fig. 4 A, B). This is consistent with work showing that the centrosome lies opposite the emerging axon in zebrafish spinal cord neurons (*14*) and Rohon-Beard cells (*63*). This positioning suggests a so-called posterior nuclear–centrosomal axis, which is also displayed by some migrating cells (*52*). Since microtubules emerge plus-end-out from the centrosome, this positioning could favour the observed focusing of plus ends towards the nascent axon (Fig. 4 C), facilitating directional transport of cellular components to facilitate axon growth. This could include other microtubules, which are slid into the nascent axon, perhaps along with their tethered neurofilaments, which we observed to accompany microtubules during protrusion emergence (Fig. 4 D). Additionally, given that neurofilaments are trafficked by microtubule motors (*37*), microtubule network polarity could also facilitate the asymmetric neurofilament deposition we observed before axon formation (Fig. 2 D–F). We found dynamic centrosome positioning to be consistent with the shifting axis of polarity during delamination, when it is positioned apically, and during axon initiation, when it is positioned in the cell body opposite the axon (Fig. 4 A). This suggests that centrosome positioning and microtubule network polarity could direct the reorientation of the polarity axis that occurs as spinal cord neurons differentiate.

Our work demonstrates a previously unreported localisation of a G protein to the intermediate filament network. This builds on a previous report that Gα protein was present in neurofilament fractions from quail (*64*). Our findings that Gαi2 accumulates on neurofilaments and colocalises with microtubules (Fig. 3) and accumulates away from the cell periphery (Fig. 1 D, G, H; Fig. S1 A) and various endomembrane structures (Fig. S1 B–D) suggests that the function of Gαi2 in this context is GPCR independent. The presence of Gαi2 at points of neurofilament–microtubule contact (Fig. 3 B) and the reduction in neurofilament–microtubule association in Gαi2-depleted cells (Fig. 5 C) suggests that Gαi2 regulates the association of these two filament types. This is supported by our finding that Gαi2 depletion reduces neurofilament, but not microtubule, accumulation in the nascent axon (Fig. 5 F, G), suggesting a functional uncoupling of these networks. The overall build-up of neurofilaments seen in Gαi2 depleted neurons (Fig. 5 D, F), together with the reduced stability of protrusions (Fig. 6 C A, C) suggests that this coupling is necessary for neurofilaments to transit away from the cell body to populate and stabilise the axon. Cross-linking structures between microtubules and neurofilaments are clearly visible in electron microscopy images, as demonstrated in *Xenopus* axons (*39*). We demonstrate close association between Gαi2 and the microtubule and neurofilament networks (Fig. 3). Thus, it is plausible that Gαi2 might also act as a linker, either alone, or as part of a larger protein complex, as it does in the context of spindle positioning in the LGN protein complex (*21*); however, further studies are required to characterise its function at the molecular level.

Gαi2 might also contribute to polarity through additional mechanisms. For example, intermediate filaments have been found to anchor signalling molecules (*65*). Attachment of Gαi2 to the polarised neurofilament network could facilitate asymmetric Gαi2 activity in the cell. This could involve local activation of polarising factors to achieve or reinforce cell polarity. For example, in migrating neural crest cells, it was recently proposed that Gαi2 regulates microtubule dynamics and activity of Rac1 (*22*), which is a key polarity regulator in neurons (*CC*). However, we did not observe an effect on microtubule polymerisation dynamics following depletion of Gαi2 (Fig. S7), suggesting that Gαi2 does not have this role in neurons. Gαi proteins are also known regulators of cell migration, notably in immune cells (*20*). Additionally, disrupted neuronal migration has been observed in Gαi2 knockdown mice (*27*). Given that cell migration involves front–rear polarisation (*67*), our findings in neurons could also apply more broadly to cell polarisation in various contexts. It will therefore be of interest to investigate the relationship between G proteins and other intermediate filaments, particularly in migrating cells.

Gαi2 is highly expressed throughout the developing vertebrate brain and spinal cord (*19, 27*). Mutations to *GNAI2*, the Gαi2 gene, in humans have been associated with intellectual disability and periventricular nodular heterotopia (*27, 68*), which are disorders associated with differences in neurite outgrowth, neuronal migration, and neuronal connectivity (*CS*). In addition, intellectual disability-associated genes converge on the cytoskeleton, particularly microtubules (*70*). Here, we show that Gαi2 regulates cell polarity and axon formation in differentiating neurons via association with the cytoskeleton. Thus, our findings offer a mechanism by which Gαi2 disruption could hinder formation of neuronal circuitry, offering potential avenues for future investigations. For example, axon instability could delay or prevent axon elongation. Defective direction of axon initiation could result in axon misdirection. Furthermore, it will be interesting to investigate potential roles for cytoskeleton-associated Gαi2 in later stages of neuronal morphogenesis, such as axon pathfinding, dendritic arborisation, or synapse formation, and in adult neurons, for example, during plasticity or regeneration.

In summary, here we uncover key features of cytoskeletal behaviour during axon formation and describe a novel role for the G protein Gαi2 in regulation of axon formation. Our results suggest that Gαi2 associates with neurofilaments, coupling them to the microtubule network, and facilitating their transit into the axon, stabilising axon outgrowth. In addition, we provide evidence that the polarised microtubule network directs polarity establishment during neuronal differentiation. These findings provide novel insights into mechanisms of cellular polarity establishment and offer an enriched view of cytoskeletal regulation during axon formation.

## Materials and methods

### Immunofluorescence and proximity ligation assays

Fertilised chicken (*Gallus gallus domesticus*) eggs were acquired from Medeggs Ltd (Fakenham, Norwich; www.medeggs.com) and incubated at 37°C to Hamburger and Hamilton (HH) stage 15–17; E3; approximately 74h of incubation). Embryos were dissected and immediately fixed in pre-warmed 4% formaldehyde solution in phosphate buffered saline (PBS) at room temperature for 2–3h, dehydrated overnight in 30% sucrose in PBS, then embedded in 1.5% Luria-Bertani (LB) agar, 30% sucrose. Embedded sections were dehydrated in 30% sucrose and frozen on dry ice. 20 μm sections were prepared with a Leica CM1950 cryostat and mounted on glass slides. Samples were permeabilised with 0.1% Triton X-100 (Sigma-Aldrich) and blocked with 1% donkey serum. Primary antibody (Table 1) incubations were performed overnight at 4°C. Incubation with phalloidin Alexa Fluor 488 (Invitrogen A12379) was carried out for 30 min at 1:500. For proximity ligation assays (PLAs), Duolink In Situ Detection Reagents (Red; Sigma-Aldrich, DUO92008 or green; Sigma-Aldrich, DUO92014) were used according to manufacturer’s instructions following primary antibody incubation as above. Samples were incubated with secondary antibodies (Table 1) for 1 h at room temperature. Where PLAs and immunofluorescence experiments were combined on the same tissue, samples were incubated with fluorescent secondary antibodies following PLA. Sections were mounted under glass coverslips using ProLong Diamond (Invitrogen, P36970) or ProLong Glass (Invitrogen, P36984) antifade mounting medium.

**Table 1:**
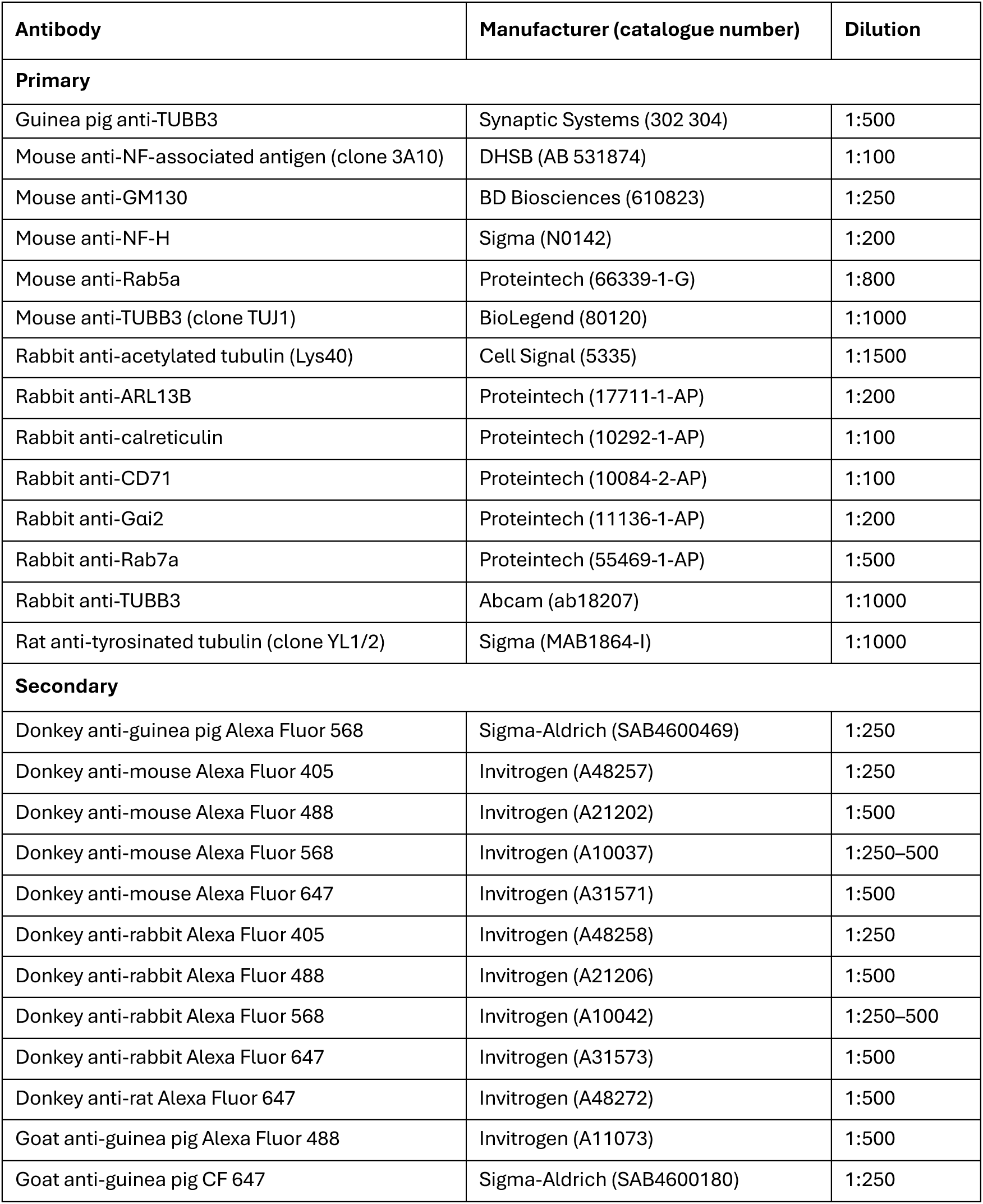
Antibodies and stains used in this study.

| Antibody | Manufacturer (catalogue number) | Dilution |
| --- | --- | --- |
| **Primary** |  |  |
| Guinea pig anti-TUBB3 | Synaptic Systems (302 304) | 1:500 |
| Mouse anti-NF-associated antigen (clone 3A10) | DHSB (AB 531874) | 1:100 |
| Mouse anti-GM130 | BD Biosciences (610823) | 1:250 |
| Mouse anti-NF-H | Sigma (N0142) | 1:200 |
| Mouse anti-Rab5a | Proteintech (66339-1-G) | 1:800 |
| Mouse anti-TUBB3 (clone TUJ1) | BioLegend (80120) | 1:1000 |
| Rabbit anti-acetylated tubulin (Lys40) | Cell Signal (5335) | 1:1500 |
| Rabbit anti-ARL13B | Proteintech (17711-1-AP) | 1:200 |
| Rabbit anti-calreticulin | Proteintech (10292-1-AP) | 1:100 |
| Rabbit anti-CD71 | Proteintech (10084-2-AP) | 1:100 |
| Rabbit anti-Gai2 | Proteintech (11136-1-AP) | 1:200 |
| Rabbit anti-Rab7a | Proteintech (55469-1-AP) | 1:500 |
| Rabbit anti-TUBB3 | Abcam (ab18207) | 1:1000 |
| Rat anti-tyrosinated tubulin (clone YL1/2) | Sigma (MAB1864-I) | 1:1000 |
| **Secondary** |  |  |
| Donkey anti-guinea pig Alexa Fluor 568 | Sigma-Aldrich (SAB4600469) | 1:250 |
| Donkey anti-mouse Alexa Fluor 405 | Invitrogen (A48257) | 1:250 |
| Donkey anti-mouse Alexa Fluor 488 | Invitrogen (A21202) | 1:500 |
| Donkey anti-mouse Alexa Fluor 568 | Invitrogen (A10037) | 1:250–500 |
| Donkey anti-mouse Alexa Fluor 647 | Invitrogen (A31571) | 1:500 |
| Donkey anti-rabbit Alexa Fluor 405 | Invitrogen (A48258) | 1:250 |
| Donkey anti-rabbit Alexa Fluor 488 | Invitrogen (A21206) | 1:500 |
| Donkey anti-rabbit Alexa Fluor 568 | Invitrogen (A10042) | 1:250–500 |
| Donkey anti-rabbit Alexa Fluor 647 | Invitrogen (A31573) | 1:500 |
| Donkey anti-rat Alexa Fluor 647 | Invitrogen (A48272) | 1:500 |
| Goat anti-guinea pig Alexa Fluor 488 | Invitrogen (A11073) | 1:500 |
| Goat anti-guinea pig CF 647 | Sigma-Aldrich (SAB4600180) | 1:250 |

Widefield images were acquired with a Zeiss Axio Cell Observer Z1 microscope equipped with an ORCA-Flash4 v2 sCMOS camera (Hamamatsu) and Colibri 7 light-emitting diode (LED) light source (Zeiss) using ZEN Blue software (Zeiss). Images were acquired with a 63x 1.4 NA (numerical aperture) oil-immersion objective (Zeiss). Z stacks were acquired with a slice interval of 0.2–0.5 μm using an exposure time of 20–100 ms. Images were deconvolved using ZEN Blue software (Zeiss) using Fast Iterative or Constrained Iterative algorithms.

Airyscan images were acquired with a Zeiss LSM 980 confocal microscope equipped with a 63x Plan-Apochromat 1.4 NA oil objective (Zeiss) and subjected to Airyscan processing. Z stacks were acquired with a slice interval of 0.2 μm.

Lattice SIM images were acquired with a Zeiss Elyra 7 SIM microscope equipped with a 63x Plan-Apochromat 1.4 NA oil objective (Zeiss) using a slice interval of 0.11 μm. Images were subjected to SIM^2^ processing in ZEN Black software (Zeiss).

STED images were acquired with a Leica TCS SP8 AOBS inverted gSTED microscope equipped with a 100x 1.4 NA HC Plan-Apochromat objective. Z stacks of 1 μm depth were acquired using a slice interval of 0.2 μm using the following settings: zoom 10, scan speed 200 Hz, line accumulation 2, STED 5%, STED 3D 5%, format 168 x 168 pixels. STED images were collected using hybrid detectors with the following detection mirror settings: Alexa Fluor 488 and Alexa Fluor 568, 0.3–6 μs gating, using the 488nm and 594nm excitation laser lines and 592 nm (5%) and 660 nm (5%) depletion laser lines respectively. Images were deconvolved in Huygens Professional software (Scientific Volume Imaging) using automated settings.

### Plasmids*, in-ovo* electroporation, and live-tissue imaging

Embryos were incubated at 37°C until HH stage 11–13 (E2, approximately 46 h incubation). Neuroepithelial cells in the spinal cord neural tube were transfected by *in-ovo* electroporation of plasmid DNA (Table 2) as detailed previously (*71*). Embryos were incubated overnight to HH stage 15–17 (E3; approximately 74 h incubation) and prepared for *ex-ovo* slice culture imaging, as previously described (*71*). Briefly, transverse sections of embryonic trunk were hand sliced and embedded in a collagen mixture (60% type I rat collagen [Corning, 354236], 1x L-15, 0.5% acetic acid, 3% sodium bicarbonate) on a glass-bottomed dish (WPI FluoroDish). Embedded slices were covered in slice culture medium (Neurobasal Medium [Gibco, 12348-017], 1x GlutaMAX [Gibco, 35050-038], 5% Optiprep Density Gradient Medium [Sigma, D1556], 1x B27 supplement, 1:1000 gentamicin) followed by recovery at 37°C, 5% CO2 for 1–3 h before timelapse imaging.

**Table 2:** Plasmids used for electroporation.

| Plasmid | Description | Concentration (ng/μL) | Source/references |
| --- | --- | --- | --- |
| dsRed-Cent2 | pCMV-dsRed-Cent2. dsRed fused to centrosomal protein Centrin 2 (human) for centrosome labelling. | 35 | Gift from Joseph Gleeson. Addgene plasmid #29523. |
| EB3-GFP | MT plus-tip protein EB3 fused to GFP for MT plus tip labelling. | 100–200 | Gift from Bill Harris. |
| NFM-eGFP | pEGFP-rNFM. EGFP fused to neurofilament protein NF-M (rat) for NF labelling. | 300 | Gift from Anthony Brown. Addgene plasmid #32907. |
| GFP-GPI | pCAGGS-GFP-GPI. GFP fused to GPI membrane anchor for membrane labelling. | 25 | Gift from Domingos Henrique, Institute of Molecular Medicine, Lisbon. |
| mKate-GPI | pCAGGS-mKate2-GPI. mKate fused to GPI membrane anchor for membrane labelling. | 50 | Gift from Kate Storey. |
| Neurog2 | pCIG-Neurog2. Neurog2 for encouraging neuronal differentiation. | 100 | Generated for this paper by University of Manchester Genome-Editing Unit. |
| PACT-TagRFP | pCMV-PACT-TagRFP. Centrosomal targeting domain PACT fused to TagRFP for centrosome labelling. | 50–100 | Gift from Kate Storey. |
| pRFPRNAi-Gai2 | Cytoplasmic RFP and microRNA operon targeting chicken Gai2 for RNAi-mediated depletion of Gai2. | 200–400 | Generated for this paper based on (47). |
| pRFPRNAi-Luc | Cytoplasmic RFP and microRNA operon targeting firefly luciferase for RNAi control. | 200–400 | (47). |

eGFP=enhanced green fluorescent protein. GFP=green fluorescent protein. GPI=glycophosphatidylinositol. IRES=internal ribosome entry sequence. Neurog2=neurogenin 2. NLS=nuclear localisation signal. RFP=red fluorescent protein. RNAi=RNA interference.

Timelapse imaging of live tissue was carried out in an enclosed environmental chamber at 37°C and 5% CO2. Images were acquired with a Zeiss Cell Observer Z1 microscope equipped with an ORCA-Flash4 v2 sCMOS camera (Hamamatsu) and Colibri 7 light-emitting diode (LED) light source (Zeiss) using ZEN Blue software (Zeiss), with an exposure time of 20 ms equipped with either a 40x 1.2 numerical aperture (NA) silicone-immersion objective (Zeiss) or a 63x 1.4 NA oil-immersion objective (Zeiss). EB3-GFP imaging for microtubule network orientation was conducted in stacks with a slice interval of 0.2–1 μm and a time interval of 1–2 s in bursts of up to 2 min for up to 24 h, allowing 1 h recovery between bursts. EB3-GFP imaging for tracking of microtubule dynamics was conducted in stacks of 10 slices, with a slice interval of 1.05–1.45 s for 1–2 min. In all other experiments, stacks were acquired with a slice interval of 1.5 μm every 10 min for up to 24 h.

### RNAi and drug treatment

Gαi2 RNAi was performed using a previously described system (*47*). Target sequences were based on the *Gallus gallus GNAI2* mRNA sequence (NCBI reference NM_205402.3) and selected using the GenScript siRNA target finder (https://www.genscript.com/siRNA_target_finder.html). The chosen target sequence for Gαi2 was 5′-AAG GAC CTC TTT GAG GAG AAG-3′. The target sequence used for the control (firefly luciferase) was 5′-TGC TGC TGG TGC CAA CCC TAT T-3′, as previously described (*47*). MicroRNA cloning site hairpins were generated by PCR using universal flanking oligonucleotides (‘hairpin forward’ and ‘hairpin reverse’) and specific oligonucleotides containing the Gαi2 target sequence (‘Gαi2 forward’ and ‘Gαi2 reverse’; Table 3). For all Gαi2 RNAi experiments, embryos were transfected by *in-ovo* electroporation at E2 after approximately 46 h incubation (HH stage 11–13) and incubated a minimum of 26–28 h (E3, approximately 74–76 h total incubation; HH stage 15–17) until fixation for immunolabelling or preparation for live imaging. Live imaging began at approximately 31 h post transfection.

**Table 3:**
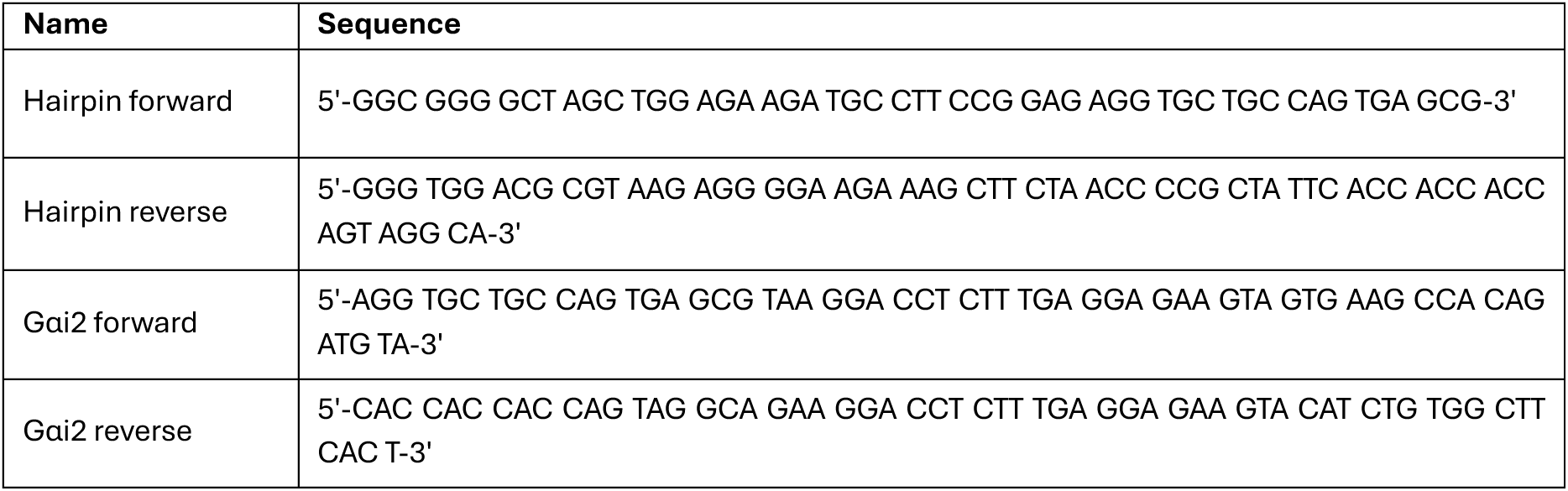
Oligonucleotides used to generate the pRFPRNAi-Gαi2 hairpin.

To depolymerise microtubules, HH stage 14–16 embryos (E3) were excised and bathed in pre-warmed and CO_2_-equilibrated slice culture medium containing either 40 μM nocodazole dissolved in dimethyl sulfoxide (DMSO) or an equal volume of DMSO alone (control). Embryos were incubated for 2 h at 37°C in 5% CO_2_ before fixation.

### Image analysis

Analysis of mean fluorescence intensity in immunofluorescence images was carried out in ImageJ or ZEN Blue software on either maximum-intensity projected z stacks or single focal planes and kept consistent within each experiment. ROIs were drawn by tracing around regions of cytoplasm (excluding the nucleus) using TUBB3 or cytoplasmic RFP as a guide. Where relevant, the cell body was divided into approximate ‘front’ and ‘rear’ regions. Given that pre-initiation cells do not have a clear cell front (ie, axon), we approximated front and rear sections of cells according to tissue anatomy. The axon is associated with the baso-ventral cell body of extending spinal cord neurons (*C*) and spinal cord neurons have previously been shown to initiate from the baso-ventral cell body in zebrafish (*14*). Cells were thus partitioned through the centre of the cell body into baso-ventral (baso-ventral cell body, plus axon in cells initiating or extending axons) and apico-dorsal (apico-dorsal cell body, plus the apical process in attached cells) regions of interest. In some cells, not all regions were intact or visible and so were omitted from measurement. For analysis of axonal Gαi2 distribution, a linear region of interest was drawn along the axon from cell body to the distal tip of the axon using the freehand line tool in ImageJ software. For growth cone analysis, growth cones of extending neurons were segmented in ImageJ software using thresholding of TUBB3. Central and peripheral regions of interest were segmented by automated erosion of a 0.5 μm region around the border and enrichment coefficients of mean gray values were calculated as above. Background fluorescence obtained from a cell-free area in the same image was subtracted from each value. In experiments where variations in staining were observed between experimental repetitions, we normalised intensities to an area of non-transfected tissue as an internal reference for staining efficiency by batch. For each batch, a scaling factor was calculated as the ratio of the global reference summary (mean across all batches) to the batch-specific reference summary (mean across all samples in a batch). All intensities in that batch were multiplied by this factor, on the basis of established ratio-based normalisation approaches used previously for imaging data (*72*). To assess regional enrichment, enrichment coefficients were calculated by dividing the value for the ROI by the mean of all ROIs per cell.

PLAs were quantified in Zen Blue by manual counting of PLA spots associated with neurofilaments.

Processing and analysis of colocalisation between channels in super-resolution images was carried out in Imaris software. 3D segmentation was performed with the Surfaces function, using thresholding to isolate and mask cells. Images were processed to remove background and smooth the signal, using the same settings for all images. Colocalisation was determined with the Coloc module using automatic thresholding. Volume overlap and Pearson’s correlation coefficient were calculated by the software. Colocalisation channels were generated in Imaris. Fluorescence intensity profile plots were generated from single focal planes in ImageJ software using the line tool.

Analysis of timelapse images was carried out in ZEN Blue (Zeiss) or Imaris (Oxford Instruments). Alongside fluorescent channels, a brightfield channel was acquired to assess tissue integrity, and analysis was restricted to the period where the tissue remained intact. Analysis of centrosome position in widefield timelapse series was carried out in 2D in ZEN Blue software (Zeiss). In the first frame that the axon was visible, the straight-line distance of the centrosome to the axon initiation site was measured and the angle of the centrosome to the axon origin site was measured with 90° directly ahead of the axon and 0° on the apical-most side of the cell.

For tracking of EB3 comet speed and distance, timelapse movies were bleach-corrected using the simple ratio algorithm (background 0) in ImageJ software and processed in Imaris software (Oxford Instruments) to improve signal-to-noise ratio. EB3 comets were tracked using the Spots function, using the automated wizard and the following settings: estimated XY diameter 0.4, background subtraction, autoregressive motion, maximum distance 0.5, and gap size 1. Quality thresholding was done manually for each cell. Tracks were filtered for duration >3 s and straightness >0.6 to reduce the false detection of tracks. Kymographs were produced in ImageJ software using the KymographBuilder plugin.

For analysis of protrusion formation, major protrusions were counted as any thick protrusion with a growth cone that grew to at least 30 μm and persisted at least 30 min. Persistence was measured as the total time a protrusion was visible before full retraction. Protrusions that were visible at the start of imaging were counted only if shorter than 30 μm when the cell was first visible. Where a cell was attached at the start of imaging, analysis began after detachment of the cell from the apical surface.

The following versions of software were used: ImageJ (FIJI), ImageJ2 v1.52–45; Imaris (Oxford Instruments), v9.8–9; ZEN Blue (Zeiss), v2.6–3.8. Unless otherwise stated, representative images were prepared in ImageJ software. Throughout, all images are oriented with the apical side of the tissue to the right.

### Statistical analysis

Statistical analyses were conducted in Prism software (GraphPad, v8.3–10.6.1). Statistical tests were selected based on normality of distribution, as determined by Shapiro-Wilk tests using an alpha of 0.05. Datasets consisting of normally distributed groups were analysed using parametric tests. Two-group datasets were analysed with t-tests (paired or unpaired). Datasets with three or more groups were analysed by one-way ANOVA, with Tukey’s multiple comparisons test for paired data or Šidák’s multiple comparisons test for unpaired data. If datasets contained any groups of non-Gaussian distribution, they were analysed by non-parametric tests. Unpaired two-group data were analysed by Mann-Whitney tests. Kruskal-Wallis tests with Dunn’s multiple comparisons tests were used for unpaired datasets with three or more groups. Analyses of multiple cellular regions between two conditions were conducted by two-way ANOVA with Šidák’s multiple comparisons test or by mixed-effects model with Geisser-Greenhouse correction for datasets where some cell regions could not be measured. An alpha of p=0.05 was used for all comparisons.

## Author contributions

Conceptualisation: RMD.

Methodology: VEH and RMD

Investigation: VEH, DA, AA, GT-T.

Formal analysis: VEH, DA, RMD.

Supervision: RMD.

Funding acquisition: RMD.

Writing – original draft: VEH and RMD.

Writing – review and editing: VEH, RMD, DA, AA, GT-T.

## Data and materials availability

All data needed to evaluate the conclusions in the paper are present in the paper and/or the Supplementary Materials. Raw data files and all reagents may be requested from the authors.

We thank our colleagues Clare Buckley, Karel Dorey, Shane Herbert and Holly Lovegrove for their comments on the manuscript. The University of Manchester Bioimaging Facility microscopes used in this study were purchased with grants from BBSRC, Wellcome and the University of Manchester Strategic Fund. We thank Peter March, Roger Meadows, and Steven Marsden from the Bioimaging Facility for facilitating imaging experiments and Chris Power and Matthew Haley from Carl Zeiss for their assistance with SIM imaging. We also thank Antony Adamson and Hayley Bennett from the University of Manchester Genome Editing Unit for their assistance with molecular cloning.

